# Belief updating in uncertain environments are differentially sensitive to reward and punishment learning: Evidence from ERP

**DOI:** 10.1101/2025.10.09.681320

**Authors:** Lingyun Xiang, Meng Liu, Weijun Li

## Abstract

Learning from rewards and punishments is crucial for adaptive decision-making, but it is still unclear how individuals integrate prediction errors to update beliefs in uncertain environments, particularly the distinctions between reward and punishment learning. We employed a probabilistic classification task and situated electroencephalography (EEG) signals within a hierarchical Bayesian framework, investigating distinctions between reward and punishment learning. Our findings indicate that participants exhibited superior learning performance in reward context compared to punishment context. Fitting the hierarchical Bayesian model revealed that punishment drives faster Bayesian belief updates, although these did not translate into improved behavioral outcomes. At the neural level, higher-level precision-weighted prediction error (pwPE2) was significantly positively correlated with FRN amplitude in the punishment context but not in the reward context, and the positive effect of pwPE2 on P300 amplitude was stronger in the punishment than the reward condition. These results provide electrophysiological signatures of punish context driving faster belief updates in uncertain environments. Our results provide novel evidence for a dual-process framework in reinforcement learning, underscoring distinct neural mechanisms underlying reward and punishment learning.

## 1 Introduction

Reward and punishment learning are two critical facets of human and animal behavior, enabling successful adaptation to environmental changes by pursuing rewards and avoiding punishments ^1^. Reinforcement learning theory provides a theoretical framework for understanding the underlying mechanisms of reward and punishment learning, suggesting that individuals extract patterns from the environment through trial-and-error and adjust their behavior accordingly, highlighting their similarities. However, prior neuroimaging studies indicate that learning from reward or punishment engages distinct neural substrates. Reward learning relies on mesolimbic dopaminergic projections, whereas punishment learning is associated with serotonergic projections^2–5^. The current study focuses on contrasting the differences between reward and punishment learning, specifically in terms of their computational and cognitive neural mechanisms. To better differentiate between reward and punishment substrates, we integrated a hierarchical Bayesian model with EEG to examine how humans implement reward and punishment learning in dynamic environments.

The Probabilistic Reversal Learning paradigm is a widely used hierarchical learning framework in learning research^6,7^, designed to simulate environments with unstable cue-outcome relationships. To maximize the probability of obtaining rewards or avoiding punishments, participants must learn the statistical associations between cues and outcomes through trial-by-trial learning. When environmental contingencies reverse, previously learned value estimates become obsolete, and the learning rate critically influences behavioral adaptation: a low learning rate may result in sluggish adjustments, while an excessively high learning rate can lead to unstable updates driven by noisy feedback ^8,9^. Consequently, this task is able to capture the impact of environmental volatility on learning behavior. Its dynamic nature makes it particularly suitable for studying the cognitive processes underlying reward and punishment learning, leading to its extensive application in both basic learning research and studies of psychiatric disorders such as autism, Parkinson’s disease, schizophrenia, and anxiety^10–13^.

With respect to Probabilistic Reversal Learning paradigm, studies of reward and punishment learning suggest that individuals exhibit marked asymmetries in task performance and learning strategies when facing reward versus punishment contexts ^10,11,14–16^. At the behavioral level, participants demonstrated different flexibility when approaching rewards and avoiding punishments. On the one hand, individuals show less flexibility in avoiding punishment than in tasks that require pursuing rewards ^15^, and have a higher learning rate under punishment tasks ^14^. On the other hand, individuals show a greater emphasis on avoiding punishment than on pursuing rewards ^16^, but their flexibility decreases instead. These opposing effects of motivational context become even more pronounced in special populations: Parkinson patients off medication exhibit profound reward–learning deficits that are ameliorated by dopamine agonists at the cost of impaired punishment learning ^10^, and adolescents with conduct disorder show selective impairments in avoiding punishment despite intact reward learning ^11^, underscoring that reward and punishment processing depend on at least partially separable neural systems. Nevertheless, purely behavioral measures cannot reveal how individuals dynamically adjust learning rates or encode prediction errors under different motivational demands; integrating computational modeling with electroencephalography (EEG) is therefore essential to uncover the temporal and mechanistic basis of these behavioral asymmetries.

Computational models capture the dynamic of learning over time—emulating a participant’s experience—and enable better mapping between behavior and neurobiology ^6,17^. In this vein, Mathys et al proposed a hierarchical Bayesian framework (Hierarchical Gaussian Filter [HGF]) ^18,19^. This model stands out for its robust explanatory power in capturing dynamic estimation of prediction errors and belief updates in uncertain environments. Its key parameters—such as learning rates (LR, controlling the speed of belief updates), precision-weighted prediction error (pwPE, quantifying the discrepancy between expected and actual outcomes), and inverse temperature parameter (ζ, reflecting decision randomness)—provide a robust framework for quantitative analysis. These parameters not only correlate with behavioral performance (e.g., accuracy) ^20,21^, but also, when combined with neuroimaging or EEG, reveal the neural underpinnings of prediction errors at different hierarchical levels ^20,22–25^. For instance, Iglesias et al utilized a probabilistic reversal learning paradigm with functional magnetic resonance imaging (fMRI) and found that low-level precision-weighted prediction errors were associated with midbrain activation (ventral tegmental area [VTA]/substantia nigra [SN]) ^23^, while high-level prediction errors correlated with activity in the septal region of the cholinergic basal forebrain, indicating the involvement of multiple neural systems in reward and punishment learning. Further, EEG studies have extended the link between HGF parameters and feedback processing. Liu et al. (2022) combined HGF with EEG and found that the P300 component independently encodes low-level pwPE_2_ (related to specific stimulus-outcome associations) and high-level pwPE_3_ (related to environmental volatility), with low-level pwPE_2_ also associated with enhanced theta-band (4-8 Hz) power, suggesting theta activity’s role in processing specific prediction errors. Similarly, Hein et al. (2021) reported that pwPE_2_ modulated centro-parietal P300 responses in reward learning tasks, further supporting that P300 is sensitive to reward-related prediction errors. Collectively, these studies demonstrate that HGF parameters not only quantify behavioral belief adjustments but also link to the multi-level characteristics of neural feedback processing, revealing that learning may rely on distinct internal mechanisms. Accordingly, this study focuses on the differential impacts of reward and punishment frameworks on learning performance, specifically examining whether systematic differences exist in behavioral outcomes and belief-updating mechanisms under volatile environments.

EEG with excellent temporal resolution is able to provide a temporal lens through the roles of feedback-related event-related potential (ERP) components within a two-stage processing model ^22,24,26–28^. The “salience-first, value-later” two-stage processing model proposed that feedback processing follows a distinct temporal structure: an early stage (∼200 ms) reflects a non-specific motivational salience signal, while a later stage (300-600 ms) encodes reward and punishment values^29^. This finding offers a theoretical framework for interpreting the roles of FRN and P300, two key feedback-related ERP components. The FRN, peaking around 200-300 ms, is the most extensively studied reward-specific ERP component, serving as a direct marker of reward prediction error (RPE) within reinforcement learning frameworks. FRN amplitude correlates with the discrepancy between expected and actual outcomes: unexpected rewards elicit a positive deflection (termed reward positivity), while unexpected losses amplify negative deflections^26,30–32^. Feedback-locked FRN responses vary based on whether expectations are confirmed or violated, with stronger amplitudes when outcomes result from self-initiated actions. Westwood and Philiastides demonstrated that an early EEG signal (∼220 ms) differentiates reward and punishment contexts, with its amplitude predicting individual differences in decision-making accuracy^21^, suggesting FRN’s role as a non-specific salience signal that facilitates subsequent behavioral adjustments. Additionally, the P300 component, occurring between 300-600 ms, is associated with attention-driven categorization of salient outcomes and the motivational significance of reward feedback, reflecting emotional and cognitive processing^33,34^. Recent studies have expanded its role to include learning and belief updating in volatile environments^22,35,36^. For instance, Hein et al demonstrated that P300 amplitude is sensitive to prediction errors in reward learning tasks^22^, suggesting a close link between this component and adaptive feedback processing. Consistent with this view, Nassar et al and Yu et al reported that feedback-locked P300 activity predicts subsequent learning behavior in changing environments^35,36^. Importantly, Nassar et al further clarified that the P300 does not merely signal unexpected outcomes but dynamically modulates learning: it facilitates adaptation when environmental changes are relevant and suppresses learning when outcomes reflect incidental noise^35^. These findings underscore P300’s role in integrating feedback for value updating and strategic adaptation. Taken together, the FRN and P300 component respectively serves to detecting early salience and promoting subsequent value updating during feedback processing. Whether the parameters of HGF model modulate the FRN and P300 components, and whether this modulate is separated between the reward and punishment contexts remain to be seen.

In summary, the current study employs a probabilistic reversal learning paradigm, combined with HGF modeling and trial-by-trial EEG analysis, to reveal the dynamic reward and punishment learning processes across behavioral, computational, and neural levels. Our primary aim was to investigate how the brain dynamically adjusts the weight of early salience detection and late value updating under different motivational contexts (seeking reward vs. avoiding punishment). We put forward the following hypotheses: (a) Behaviorally, punishment contexts may decrease behavioral flexibility and enhance learning rates and prediction errors in the HGF framework; (b) Neurally, HGF parameters may differentially modulate FRN and P300 under reward and punishment contexts respectively, reflecting the distinct neural mechanisms underlying reward and punishment learning. Specifically, the parameter modulation on FRN amplitude (i.e., detecting early salience) would be stronger in the punishment than the reward context, reflecting enhancing sensitivity to feedback in punishment context. Additionally, the parameter modulation on P300 amplitude (i.e., value updating) would be more prominent in the punishment than the reward context, reflecting faster belief updates in the punishment context.

## 2 Results

### 2.1 Behavioral results

#### 2.1.1 Accuracy

The fitting results of the GLMM regarding the Accuracy are shown in Table S1 (see supplementary material). Participants exhibited superior performance in the reward context compared to the punishment context: under stable conditions, accuracy was markedly lower in the punishment context (*β* = −0.22, *SE* = 0.05, *z* = −4.20, *p* < .001) relative to the reward context. Environmental volatility also impaired performance, with accuracy significantly reduced in volatile environments within the reward context (*β* = −0.52, *SE* = 0.05, *z* = −10.43, *p* < .001). Critically, a significant interaction between context and environment type emerged (*β* = 0.31, *SE* = 0.07, *z* = 4.39, *p* < .001), indicating that the detrimental impact of volatility on accuracy was more pronounced in the reward context than in the punishment context, where punishment appeared to partially mitigate volatility-induced declines—highlighting a key asymmetry in how reward and punishment contexts modulate adaptation to uncertainty (see Figure 1A).

**Figure 1.**
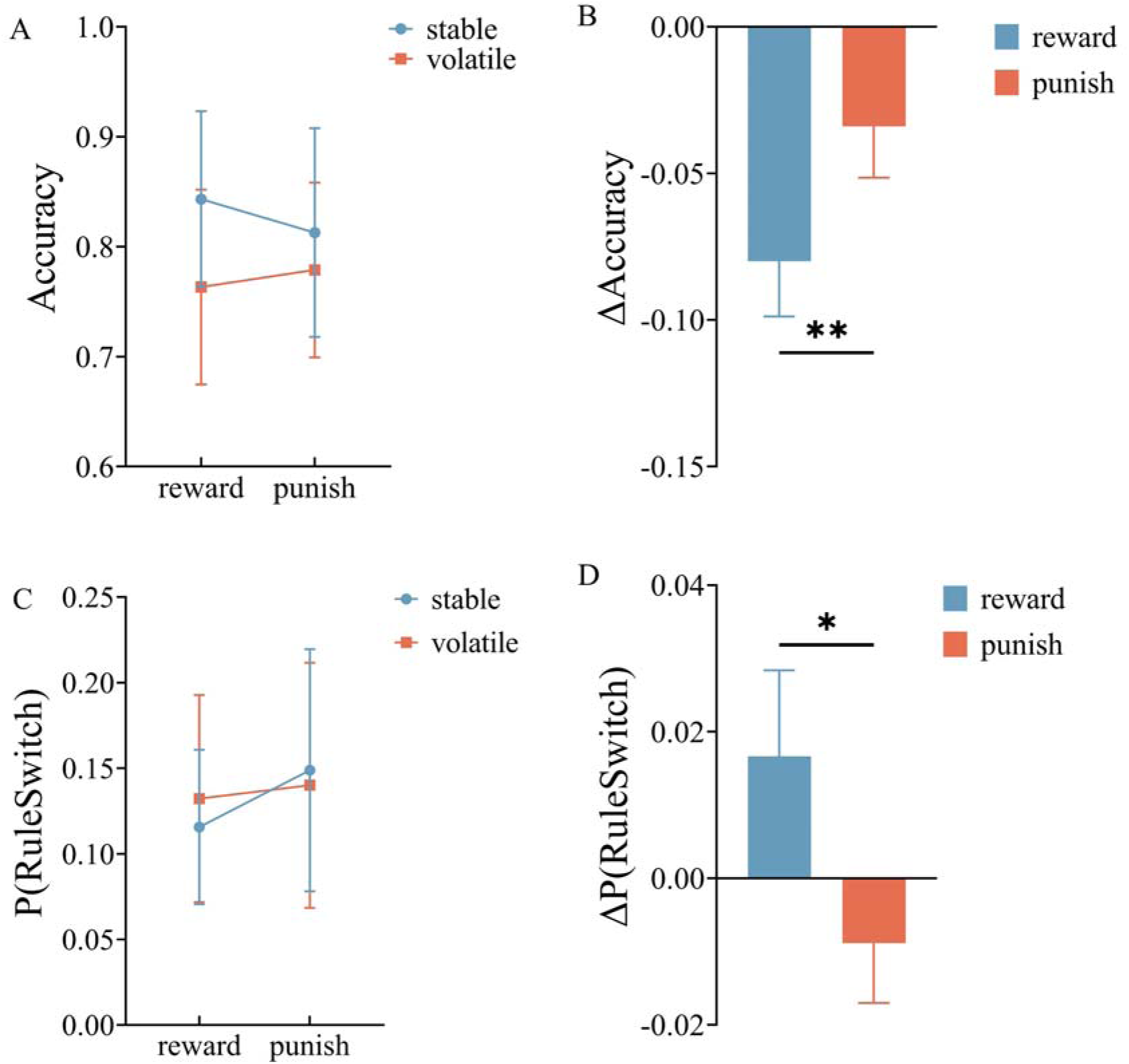
Behavioral results. (A) GLMM analysis of accuracy revealed a significant interaction between environmental type and context. (B) Paired-sample t-tests indicated a significant difference in accuracy between reward and punishment contexts. (C) GLMM analysis of post-task switching rate *P*(RuleSwitch) revealed a significant interaction between environmental type and context. (D) Paired-sample t-tests indicated a significant difference in *P*(RuleSwitch) between reward and punishment contexts. \**p* < 0.05, \*\**p* < 0.01

For completeness, we analyzed ΔAccuracy using the paired *t* test. The analysis results show that, ΔAccuracy values were negative in both conditions, with ΔAccuracy in the reward context (*M* = -0.08 ± 0.02) being significantly lower than in the punishment context (*M* = -0.03 ± 0.02), *t* (34) = -3.01, *p* = 0.005 (see Figure 1B). This pattern suggests that participants were less flexible in avoiding punishment than in pursuing reward, highlighting an asymmetry in adaptive adjustments under different motivational contexts.

#### 2.1.2 *P*(RuleSwitch)

GLMM fitting results for the probability of switching rules after error feedback (*P*(RuleSwitch)) are presented in Table S2. Participants showed a lower propensity to switch in the reward context compared to the punishment context, as evidenced by a significant main effect of task type: under stable conditions, the probability of switching was markedly higher in the punishment context (β = 0.3, *SE* = 0.06, *z* = 5.11, *p* < .001) relative to the reward context. Environmental volatility also increased switching, with the probability significantly elevated in volatile environments within the reward context (β = 0.16, *SE* = 0.06, *z* = 2.63, *p* = .008). Critically, a significant interaction between task type and environment type emerged (β = -0.23, *SE* = 0.08, *z* = -2.82, *p* = .005), indicating that the facilitative effect of volatility on switching was attenuated in the punishment context relative to the reward context, where punishment appeared to reduce excessive exploratory behavior in uncertain environments (see Figure 3C).

Similarly, we used the paired sample t-test to analyze Δ*P*(RuleSwitch). The results showed that Δ*P*(RuleSwitch) in the reward context (*M* = 0.017 ± 0.01) was significantly higher than in the punishment context (*M* = -0.01 ± 0.01), *t* (34) = 2.41, *p* = 0.022 (see Figure 3D). This finding further supports the notion that participants exhibited reduced flexibility in adapting their behavior to avoid punishment compared to pursuing reward.

### 2.2 Model Fitting

#### 2.2.1 Model comparison

We fitted each model using response data from 34 participants and conducted a group-level comparison of the three models using random-effects Bayesian model selection (BMS). The log-model evidence (LME) values obtained from model fitting were used as random effects, implemented via the SPM12 toolkit (https://www.fil.ion.ucl.ac.uk/spm/). This approach estimated the expected posterior probabilities (EXP_P), representing the probability that a randomly selected participant’s data is best explained by a given model, as well as the exceedance probability (XP) and protected exceedance probability (PXP), which indicate the probability that a given model outperforms all other models in the model space (see Table 1). The results demonstrated that the HGF model outperformed the other two models, and thus, we selected the HGF model as the optimal model for subsequent analyses.

**Table 1.** Model comparison.

|  | HGF | RW | SK1 |
| --- | --- | --- | --- |
| EXP_P | 0.98 | 0.01 | 0.01 |
| XP | 1 | 0 | 0 |
| PXP | 1 | 0 | 0 |

#### 2.2.2 Model parameters

For the probabilistic learning rate (learningrate2, LR2), the absolute value of the precision-weighted prediction error (|pwPE2|) and ζ, we conducted paired *t* tests. For LR2, the difference between the reward context (*M* = 0.937 ± 0.29) and the punishment context (*M* = 1.148 ± 0.62) was significant, *t*(33) = -2.35, *p* = .025, the learning rate was higher in the punishment context. For |pwPE2|, the probabilistic prediction error in the punishment context (*M* = 0.48 ± 0.27) was significantly higher than in the reward context (*M* = 0.39 ± 0.12), *t*(33) = -2.41, *p* = .022. For the parameter ζ, the difference between the reward context (*M* = 2.43 ± 1.57) and the punishment context (*M* = 2.18 ± 0.97) was not significant, *t*(33) = 1.13, *p* = .27 (see Figure 2).

**Figure 2.**
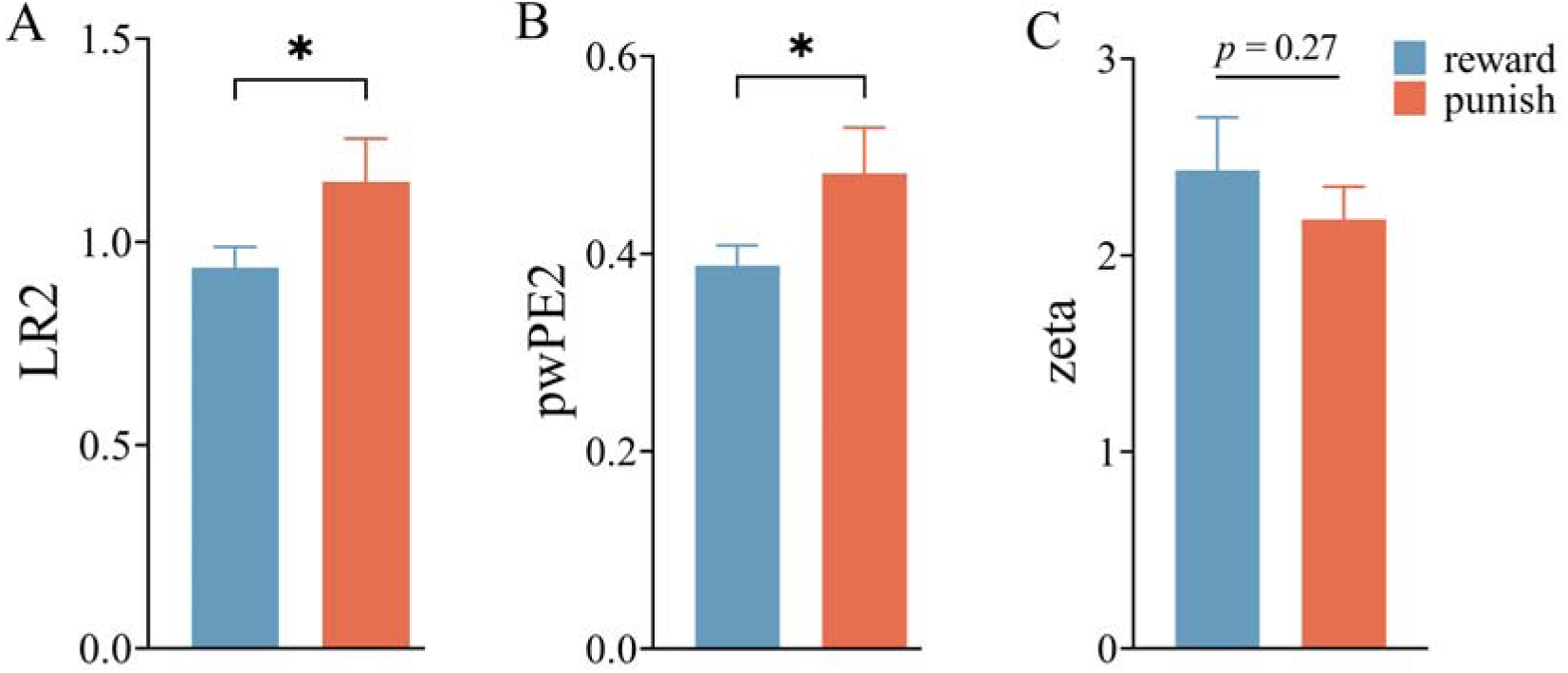
Model parameters results. (A) Probabilistic learning rate (LR2). (B) Probabilistic prediction error (|pwPE2|). (C) Inverse temperature parameter (ζ). \**p* < 0.05

### 2.3 Combined analysis of EEG and model parameters

We compared the results of models with and without parameter modulation, incorporating Context and Environmental volatility (with parameter modulation including the standardized |pwPE2| model parameter in the LMM) as fixed factors, setting reward and stable conditions as the baseline. A linear mixed-effects model with participants as random intercepts was used, with trial-by-trial data from the electrodes and time windows selected in the original analysis serving as the dependent variable. By comparing the linear mixed models with and without parameters under the two components, we found that the model with parameters was always superior to the one without parameters, which indicates the importance of this parameter for the electroencephalogram component (see Table S3).

#### 2.3.1 Parameter-free modulation

The fixed effects results for the FRN without parameter modulation are presented in Table S4. Compared to the stable environment, the volatile environment significantly reduced FRN amplitude (β = -0.75, *SE* = 0.21, *t* = -3.64, *p* <.001), indicating that the volatile environment elicited a more negative FRN compared to the stable environment.

The fixed effects results for the P300 without parameter modulation are presented in Table S5. The P300 amplitude in the volatile environment was significantly reduced compared to the stable environment (β = -1.12, *SE* = 0.20, *t* = -5.57, *p* <.001), indicating that the stable environment elicited a more positive P300 amplitude.

#### 2.3.2 Parameter Modulation

We evaluated the non-modulated model against the model including pwPE2 using a likelihood ratio test, which revealed that the model with the added parameter significantly outperformed the non-modulated model.

For the FRN component, the fixed effects results with the standardized |pwPE2| included as a fixed factor are presented in Table S6. Beyond the effect of environment type, a significant interaction was found between Context and |pwPE2| (β = 0.9, *SE* = 0.21, *t* = 4.38, *p* <.001). However, in the reward context, the main effect of |pwPE2| yielded a β value of only -0.06, indicating that |pwPE2| did not significantly influence FRN amplitude in the reward context. In contrast, in the punishment context, |pwPE2| was significantly positively correlated with FRN amplitude (see Figure 3).

**Figure 3.**
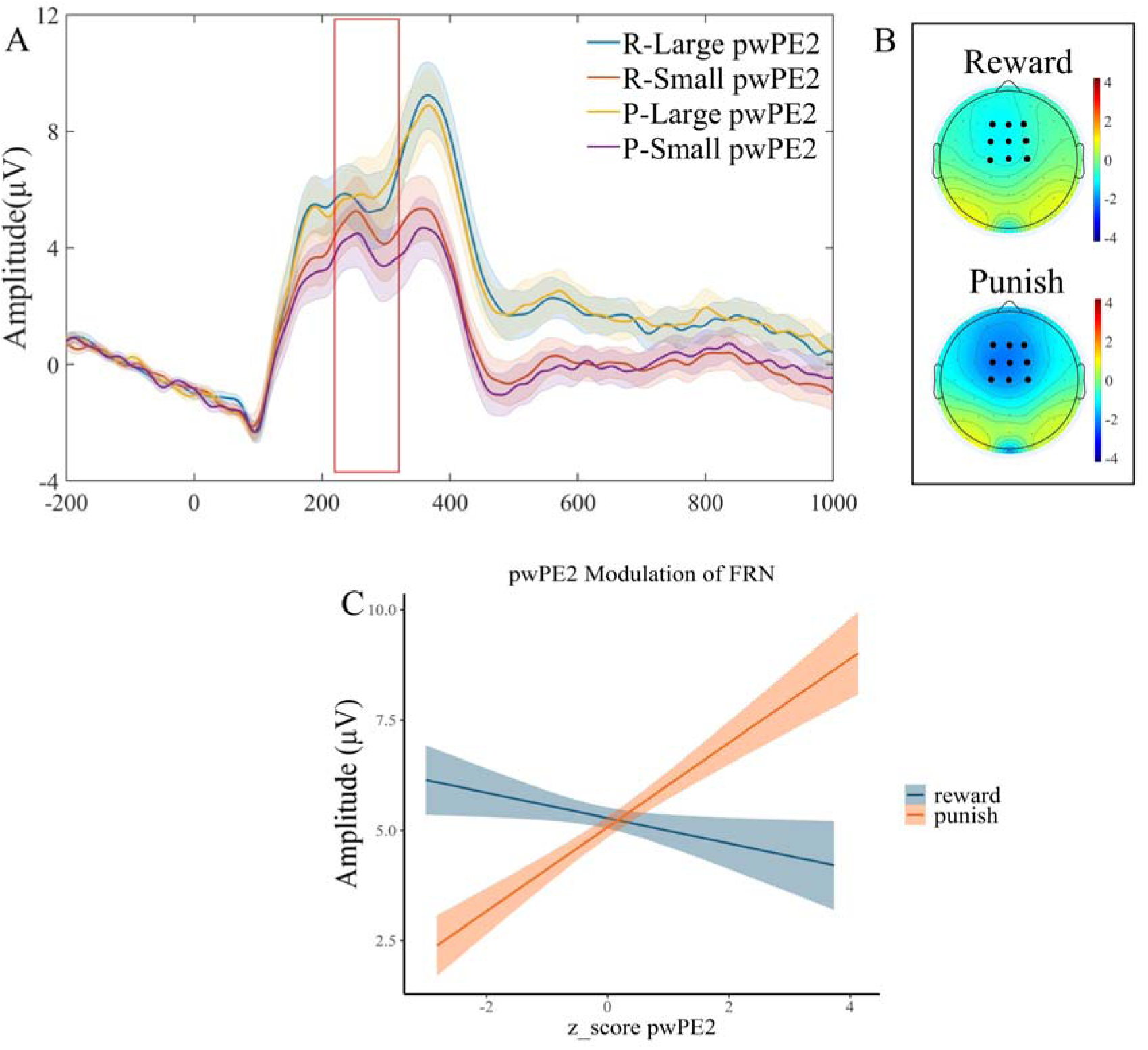
The results of the combined analysis of FRN and model parameters. (A) FRN waveform plot. (B) topographic maps showing the difference between low and high |pwPE2| for reward (top) and punishment (bottle) conditions. (C) Interaction effect of context × parameter on FRN amplitude under parameter modulation, with shaded areas representing the 95% confidence interval (CI).

The fixed effects results for the P300 component with the inclusion of the standardized absolute precision-weighted prediction error (|pwPE2|) are presented in Table S7. In addition to the effect of environment type, we observed a significant main effect of |pwPE2| (β = 1.50, *SE* = 0.15, *t* = 9.81, *p* <.001) and a significant interaction between Context and |pwPE2| (β = 0.54, *SE* = 0.20, *t* = 2.73, *p* = .006). The main effect of |pwPE2| indicates that in the reward context, the P300 effect increased with larger |pwPE2| values. The significant interaction between Context and |pwPE2| suggests that the positive effect of |pwPE2| on P300 amplitude was stronger in the punishment context compared to the reward context (see Figure 4).

**Figure 4.**
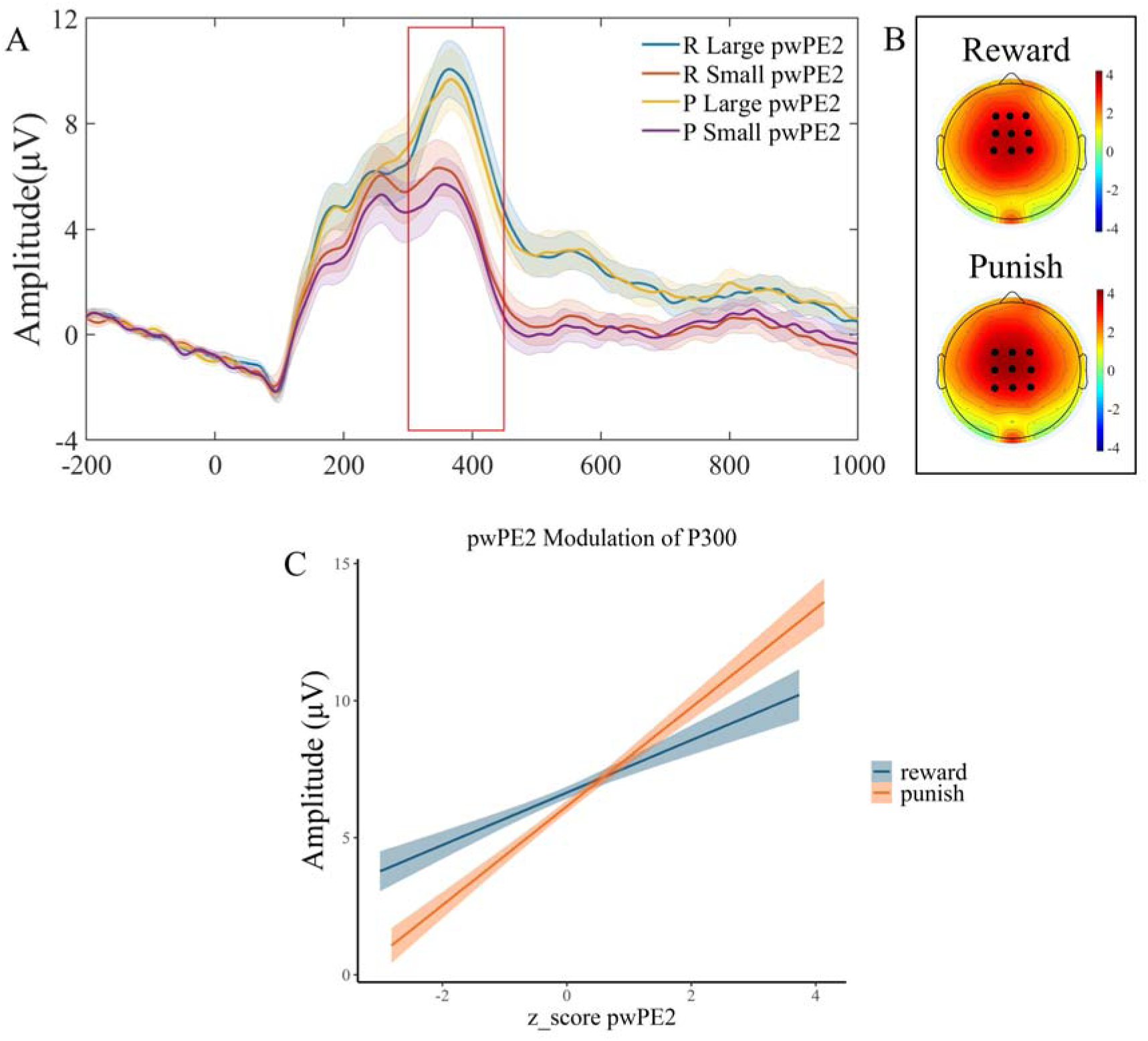
The results of the combined analysis of P300 and model parameters. (A) P300 waveform plot. (B) topographic maps showing the difference between low and high |pwPE2| for reward (left) and punishment (right) conditions. (C) Interaction effect of context × parameter on P300 amplitude under parameter modulation, with shaded areas representing the 95% confidence interval (CI).

## 3 Discussion

This study combined a probabilistic reversal learning task with trial-by-trial EEG and Hierarchical Gaussian Filter (HGF) modeling to examine the cognitive mechanisms of reward and punishment learning under volatility. We found that the punishment context exhibited a higher probability of switching rules after negative feedback in stable environments, with this effect attenuated in volatile environments. The HGF revealed higher learning rates and larger precision-weighted prediction errors in punishment contexts, indicating stronger responses to uncertainty. Analysis of the modulatory effects of the |pwPE2| on EEG signals showed that FRN was selectively enhanced by punishment feedback, and P300 scaled positively with |pwPE2|. Critically, |pwPE2| modulated FRN only in the punishment context, while it positively modulated P300 in both contexts, with a stronger effect for punishment. These results reveal a functional dissociation between early error signals and late belief-updating signals in reward and punishment learning. Besides, our results underscore that the commonly presumed advantage of reward-based learning is not universal but depends on environmental uncertainty.

Behaviorally, the reward advantage was only reflected in the stable environment; under volatility, performance did not differ between reward and punishment. In addition, the negative effect of volatility on accuracy was larger in the reward context. On the one hand, our results are consistent with those of previous studies^24,37–39^, and on the other hand, we extend the conclusions of previous studies. Previous studies reported better learning in reward than punishment contexts ^38^ and longer time to avoid punishment ^39^. However, our results showed that when the environment shifted from stable to volatile in reward contexts, task performance markedly declined, highlighting the significant impact of environmental volatility on learning. But the interaction between Context and Environmental volatility revealed that the negative effect of volatility on performance was partially mitigated in punishment contexts, showing a weaker detrimental effect compared to reward contexts. A similar pattern was observed for the probability of switching rules after error feedback: switching was least frequent in the reward-stable condition, while it significantly increased in the punishment-stable condition, consistent with the possibility that punishment inflates perceived volatility and promotes over-exploratory switching. Functionally, this may show that reward contexts appear more vulnerable to volatility in terms of accuracy, while punishment contexts seem to be less affected but at the price of under-adjusting exploration to the true level of environmental change.

Reduced behavioral flexibility in the punishment context further reveals the differences across the reward-punish contexts. Our results indicate that participants increased exploratory switching and accepted short-term costs when pursuing rewards, but did not adjust switching proportionally when avoiding punishment. In other words, avoidance framing dampened flexibility: participants did not sufficiently increase rule switching as volatility rose and thus maintained a relatively rigid response policy. This rigidity can superficially appear “resistant” to volatility in accuracy terms (a smaller drop in punishment), yet it reflects under-adaptation rather than robustness. These observations are consistent with prior reports of reduced flexibility under aversive motivation ^15,40^. This asymmetry may indicate that aversive motivational states may lead to decreased adaptive control, with individuals developing relatively higher tolerance for errors in order to be able to reduce potentially fatal error production in the context of avoiding punishment ^15,41^.

Our computational modeling results suggest that reduced behavioral flexibility in the punishment context is underpinned by significant differences in the computational learning mechanisms. That is, learning rates were higher in punishment contexts compared to reward contexts, and similarly, higher in volatile environments compared to stable ones ^42^. Individuals exhibited faster belief updating in negative contexts, and assigned greater weight to prediction errors in unstable environments. These findings align with Bayesian learning theory, where learners update beliefs by inferring environmental volatility^18,19^. The elevated punishment-context learning rate accords with neurobiological models emphasizing ACC-mediated error monitoring and avoidance learning ^26^. Additionally, increased |pwPE2| in volatile environments supports a core assumption of hierarchical learning models, namely that individuals adjust learning strategies based on perceived uncertainty. Contrary to frameworks that emphasize reward-driven learning ^3^, the present pattern suggests that threat avoidance may have an evolutionary priority over reward pursuit under uncertainty ^43^.

However, we did not find any differences between the reward and punishment in the inverse temperature parameter (ζ). Within the decision model, higher ζ yields more deterministic, policy-consistent choices for a given belief state, whereas lower ζ permits more exploratory or noisy responding. Notably, applying high-definition theta-frequency tACS over right DLPFC during probabilistic reversal learning increased low-level learning rate and uncertainty and amplified low-level pwPEs, yet left the decision model’s temperature parameter unchanged—implicating frontal theta in belief updating rather than in altering choice stochasticity per se ^25^. This dissociation suggests that motivational framing primarily reconfigured perceptual learning (how beliefs are updated about cue–outcome probabilities) rather than the determinism of the choice rule ^44^. These computational findings provide a bridge to subsequent EEG analyses, suggesting that behavioral differences in reward and punishment learning may stem from distinct neural processing priorities at different stages.

The present model-based ERP results revealed that the |pwPE2| significantly and positively modulated FRN amplitude only in the punishment context framework, with no significant effect on FRN in the reward context. Previous EEG studies have demonstrated that the FRN is considered an early, rapid encoding of “unexpected outcomes” or “negative prediction errors” ^26,30^. Our finding aligns with the notion that the increased engagement of ACC-mediated rapid error detection when precision-weighted errors associated with negative outcomes grow, producing a larger FRN deflection^26,30^. Intracranial evidence further shows anatomically dissociable prediction-error signals for reward and punishment^39^, supporting partly segregated evaluative channels. By contrast, although |pwPE2| also quantifies error magnitude under reward, it did not modulate FRN—suggesting that early error-monitoring mechanisms are preferentially tuned to aversive outcomes ^26^. In short, FRN selectively encodes punishment prediction errors.

Furthermore, our model-based ERP results also demonstrated that the P300 was not only positively modulated by the main effect of the precision-weighted prediction error (|pwPE2|) but also exhibited stronger modulation in the punishment context compared to the reward context. Previous EEG studies reported that the P300 is widely regarded as a later cognitive processing marker of “environmental salience” or “demand for belief updating”^33,34^. This ERP amplitude acts as a late feedback stage that integrates unexpected feedback into belief updating while being modulated by motivational salience ^29^. Larger prediction errors increased P300 across contexts, reflecting a general attentional response, but the effect was amplified under punishment, likely due to the higher motivational and emotional salience of unexpected outcomes in aversive contexts, which recruit greater cognitive resources. Prior work has tied the centro-parietal P300 to adjustments in learning rate and belief revision, with its relationship to trial-wise updating depending on whether surprise signals a true change point or noise ^35^. Multimodal EEG-fMRI studies likewise point to separable yet interacting value systems, with later parietal signals reflecting outcome evaluation beyond initial salience detection ^29^. Within hierarchical Bayesian formulations, P300 has been linked to precision-weighted error terms that drive belief updating, a mapping supported by recent EEG-HGF work showing trial-wise associations between pwPEs and late potentials ^24^ and causal modulation of low-level learning dynamics—without changing decision temperature—via theta-frequency stimulation of control circuitry^25^. In this framework, |pwPE2| reflects belief updates under uncertainty, and the enhancement of P300 may correspond to this belief adjustment process, particularly in punishment where motivational salience is higher ^21^.

Collectively, our results showed both FRN and P300 components tracked trial-wise precision-weighted errors, indicating a shared computational substrate—a hierarchical updating process—expressed at distinct timescales. FRN rapidly detects the salience of unexpected outcomes, while P300 integrates feedback to update beliefs ^29^. Critically, the modulation patterns across these stages differ between reward and punishment contexts. FRN is specifically modulated by |pwPE2| in the punishment context, reflecting the early system’s prioritized response to negative feedback. In contrast, P300 is modulated in both reward and punishment contexts, but the effect is stronger in the punishment context, indicating that late value updating requires greater cognitive resources in aversive contexts. Prior work aligns with this division of labor: early fronto-central signals encode outcome-related salience/prediction error, while later centro-parietal activity indexes the incorporation of those errors into belief states and subsequent learning, with the mapping between EEG and learning itself being conditional on environmental statistics ^29,35^. Within HGF-based analyses, pwPEs have been shown to relate both to oscillatory markers and to late evoked activity that scale with hierarchical uncertainty, and causal perturbation of frontal theta selectively shifts learning parameters while sparing decision temperature—supporting a two-stage ^24,25^, learning-centric account that our results extend to a reward-versus-punishment comparison. Altogether, early salience detection (FRN) provides a foundation for late value updating (P300) in the punishment context, reflecting a hierarchical progression between stages. However, in the reward context this early salience processing is blunted, with greater reliance on the late stage for integrating long-term expectations.

## 4 Method

### 4.1 Participants

Based on previous studies and following standard recommendations ^25,45^, an a priori power analysis was conducted using G*Power 3.1.9.2 ^46^ to determine the required sample size. Given a repeated-measures within-subjects design with two factors (2 × 2), we estimated the required sample size to detect a medium effect size (Cohen’s *d* = 0.5), with a statistical power of 0.80 and an alpha level of 0.05. The analysis indicated that a minimum of 34 participants was required to achieve adequate power. Thirty-six university students participated in this experiment. All participants had normal or corrected-to-normal vision, no hearing or neurological impairments, and no color blindness or color weakness. The study was approved by the Research Ethics Committee of Liaoning Normal University (**No.LL2025229)**, and all subjects gave written informed consent in accordance with the Declaration of Helsinki before participation. One participant did not complete the experiment, and another was excluded due to excessive EEG artifacts. Thus, the final sample consisted of 34 participants (16 males), aged 19-25 years (*M* = 22.3, *SD* = 2.56).

### 4.2 Materials

In probabilistic classification task, the classification stimuli were eight images selected from the image library provided by Kool et al. (2010). This image library has been used as a classification stimulus in many probabilistic classification tasks ^15,25^. In the current study, images were randomly labeled as S1, S2, …, S8 for each participant, constituting 4 pairs of paired images with different colors. To avoid the mutual influence of learning at different content stages and prevent participant fatigue during the experiment, different pairs of images were presented in each block. In each trial, they were shown one of two images and were required to guess whether it belonged to Category A or Category B, with the probabilities of the two images belonging to a given category being complementary.

### 4.3 Procedure

We employed a within-subject 2 (Context: reward, punishment) × 2 (Environmental volatility: stable, volatile) factorial design. On each of four task blocks, participants completed a probabilistic categorization task based on previous studies^10,22,24,38^. The task was programmed in PsychoPy (version 2024.2.4) ^48^ and run on a PC; participants were seated upright in front of a 1920 × 1080 monitor with a 60 Hz refresh rate and stimuli were presented centrally. On each trial one image from a paired stimulus set was displayed for 1,200 ms, during which participants were required to classify the stimulus as category A or B by pressing the corresponding response key (response window = 1,200 ms). Response-key mappings were balanced across trials to minimize practice-related biases. After the stimulus epoch a blank screen was presented for a jittered interval of 800-1,200 ms, after which trial-wise feedback was shown; following feedback a fixation cross appeared for another jittered 800–1,200 ms before the next trial began (see Figure 5A). Feedback valence and format differed between the reward and punishment contexts ^38^: in the reward context a correct response produced positive feedback displayed in green (“+25 points”) while an incorrect response produced neutral feedback in red (“+0 points”); in the punishment context a correct response produced no point loss (displayed as “-0” in green) whereas an incorrect response produced negative feedback in red (“-25 points”) (see Figure 5B). Participants were informed prior to the main experiment that their remuneration consisted of a fixed participation fee (RMB 35) plus a performance-dependent bonus derived from accumulated points, converted at a rate of 100 points = 1 RMB.

**Figure 5.**
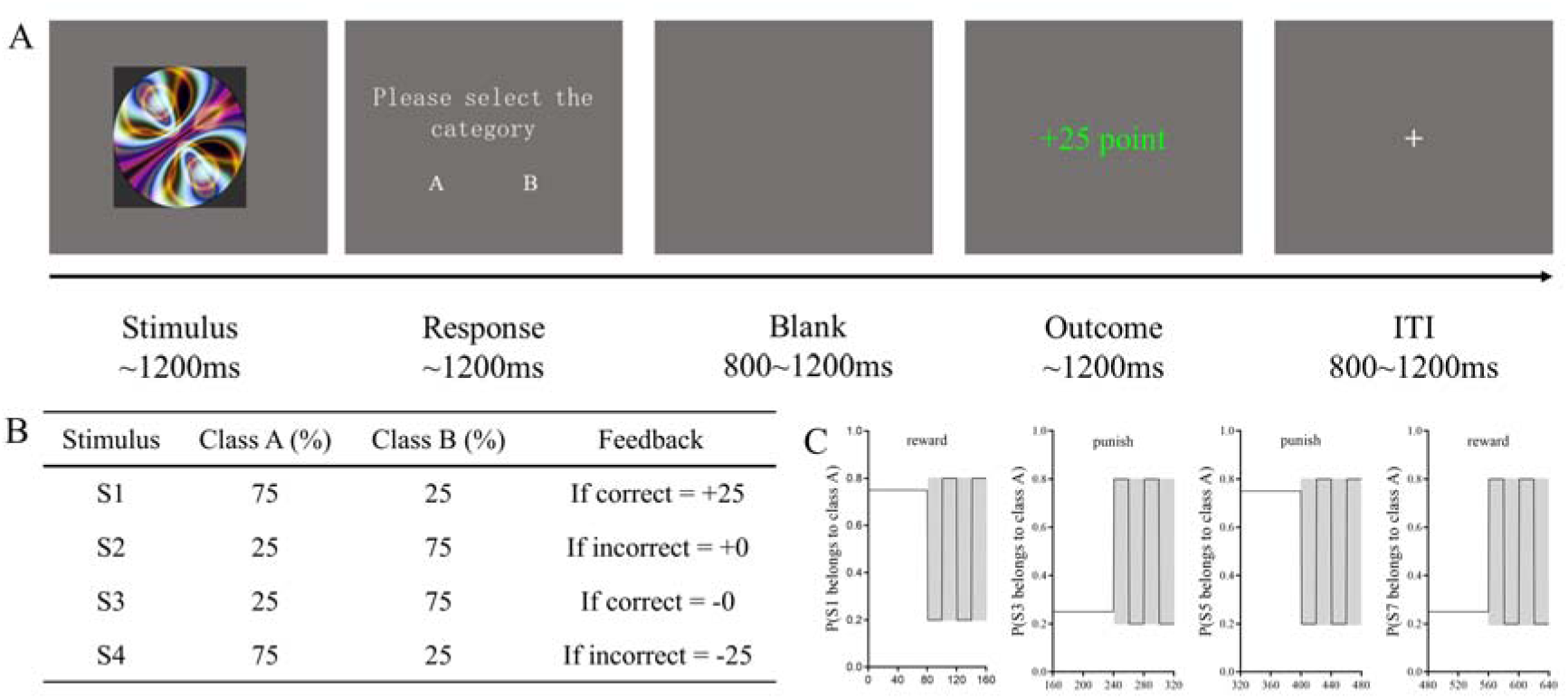
Experimental design. (A) Flowchart of a single trial. (B) Point settings in reward and punishment contexts: In the reward context, correct responses add 25 points, while incorrect responses add 0 points; in the punishment context, correct responses subtract 0 points, while incorrect responses subtract 25 points. (C) Cue-outcome contingency of images over time in the experiment. In the first 160 trials, images S1 and S2 were presented. In the stable environment, the probability of S1 belonging to Category A was fixed (0.75/0.25), whereas in the volatile environment, the probability shifted to 0.8/0.2 and reversed four times. The remaining trials followed a similar structure.

In total, each participant completed two reward and two punishment blocks; each block comprised 160 trials. Block order was counterbalanced across participants such that subjects were randomly assigned to a reward-first or punishment-first sequence, while within each block the two environmental conditions (stable, volatile) were implemented according to the task structure. In the stable environment the probability that the paired images belonged to the dominant category was held at 0.75 (non-dominant = 0.25); in the volatile environment cue–outcome associative strength varied over time, with category probabilities switching between 0.2 and 0.8 on four occasions (see Figure 5C). To avoid excessive difficulty and carry-over effects associated with beginning in a volatile context, each block was presented with the stable environment first. Prior to the main experiment participants completed a practice session that included both reward and punishment contexts (20 trials per context) and contained one category reversal so that the practice mirrored the structure of the experimental blocks; the main experiment proceeded after participants confirmed they understood the task. The entire experiment lasted approximately 1hour, with participants taking a brief rest every 60 trials.

### 4.4 Hierarchical Gaussian Filter

We employed the Hierarchical Gaussian Filter (HGF) model developed by Mathys et al^18,19^ to estimate each participant’s learning process and belief trajectories. The three-layer model, depicted in Figure 6, describes the states at each level. The first-layer state, X1, represents the binary variable of the experimental stimuli: in trial (k), either S1 = A & S2 = B (X1^(k)^ = 1) or S1=B & S2=A(X1^(k)^=0). The second-layer state, X2, describes the probability of reward occurrence. The third layer, X3, represents the log-volatility of the second layer. These hidden states (X2 and X3) are assumed to evolve via a Gaussian random walk, with their mean at trial k-1 centered on the previous value and variance dependent on the state of the layer above. The learning parameters are ω2, ω3 and ζ. Consistent with prior studies^22,24^, we fixed κ at 1.

**Figure 6.**
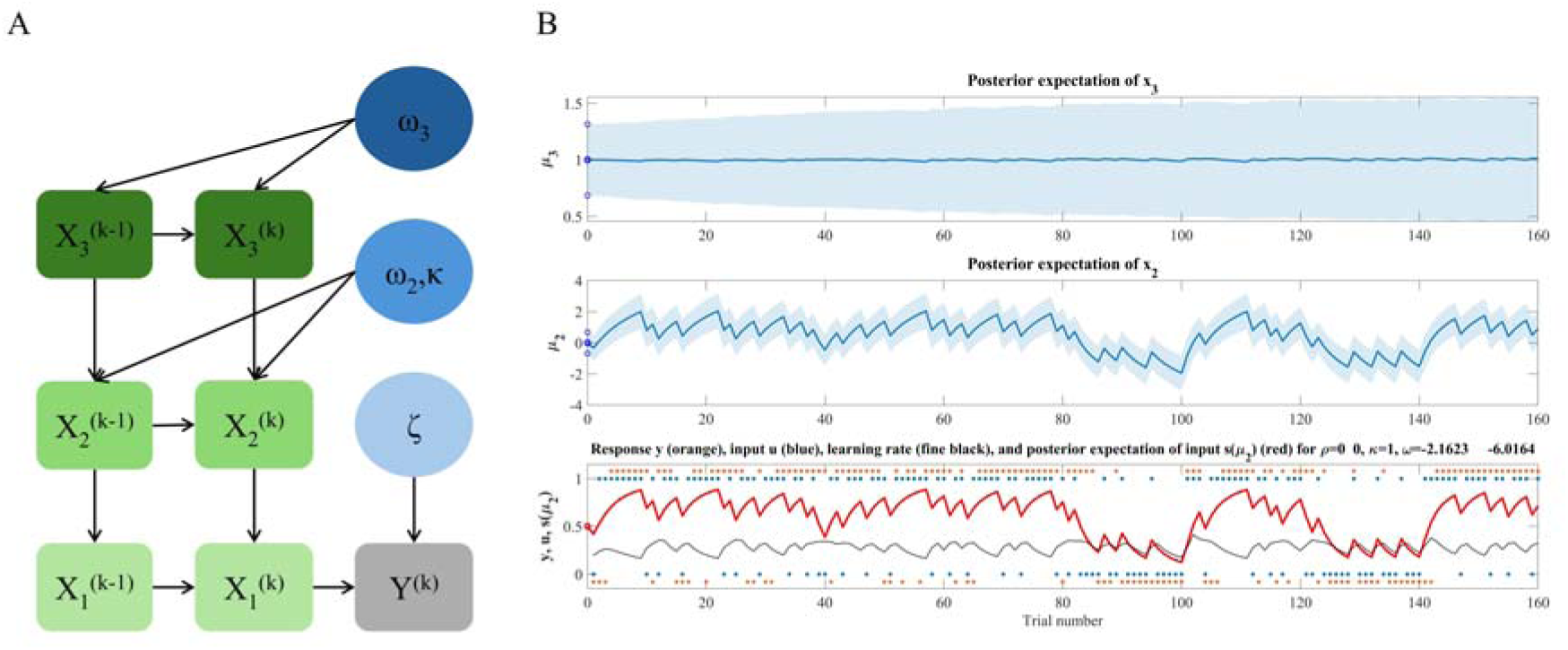
Schematic diagram of the three-level HGF model. (A). with κ fixed at 1. Typical results of model fitting for a single participant (B). At the lowest level, u (blue dots) represents the cue-outcome association for each trial, and y (orange dots) represents the participant’s response (1for S1 = A & S2 = B, 0 for S1=B&S2=A). The red line, s(μ2), indicates the participant’s estimated cue-outcome tendency *x*2, calculated using a sigmoid function. The learning rate for the cue-outcome association at the lowest level is depicted by a black line. At the second level, μ2 (green line) represents the participant’s estimate of the cue-outcome tendency *x*2. At the highest level, μ3 (blue line) represents the estimate of volatility *x*3.

The update equations for the HGF for each trial and level i (i = 2, 3) are as follows:

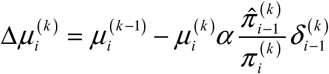

The posterior mean update, 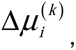 represents the difference between the posterior expectations of the previous trial k-1 and the current trial k. The lower-level prediction error, 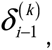, is used to update the belief at the current level. The precision ratio,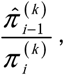 is defined where 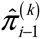 is the expected precision of the lower level, and 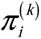 is the expected precision of the current level. Precision,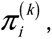 is defined as the inverse of the variance of the posterior expectation:

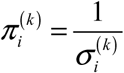

In the HGF model, the precision-weighted prediction error (pwPE) serves as the core driving signal, calculated as the product of the prediction error magnitude and its associated precision. This acts as a “correction signal” used by the brain to update beliefs, widely regarded as a critical neural computational mechanism in perception and learning.

The HGF model employs a unit-square sigmoid observation model to link the posterior estimates of beliefs (what the participant thinks is likely) to the expressed decisions y(k). In this model, the predicted probability m(k) that the presented image belongs to a specific category (e.g., S1=A & S2=B) on trial k is linked to trial-wise category predictions via the softmax (logistic sigmoid) function:

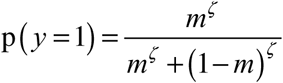

where ζ is a subject-specific temperature parameter, interpreted as the inverse of decision noise. This parameter is the only free parameter in the unit-square sigmoid observation model and serves to capture the degree of deterministic association between y and m. Higher ζ values indicate that participants are more likely to choose the option that is more congruent with their current belief. For a more detailed understanding of the specific equations and construction of the HGF model, refer to the works of Mathys et al^18,19^.

### 4.5 Model space

We evaluated three computational models of learning. The first was a three-layer HGF model, which integrates key principles of reinforcement learning and Bayesian learning. The other two control models were the Rescorla-Wagner (RW) model and the Sutton K1 (SK1) model. The RW model, one of the most widely used and simplest reinforcement learning models, assumes that expected errors drive belief updates under a fixed learning rate. The SK1 model incorporates an adaptive learning rate but lacks a hierarchical structure. Comparing these two models with the HGF model allows us to assess whether participants engaged in hierarchical learning and adaptively updated their learning processes. Model comparison was conducted using the protected exceedance probability metric from random-effects Bayesian model selection (BMS)^49^.

### 4.6 EEG recording and preprocessing

EEG was recorded using a 64-channel Brain products setup, and all electrodes were arranged in accordance with the extended international 10-20 system. The impedance for all electrodes was checked and reduced until it was below 5 kΩ. The FCz electrode was used as the reference electrode while recording, and a 100-Hz low-pass filter was applied. The average value of TP9 and TP10 amplitudes of the left and right mastoid process electrodes was used for re-reference during offline analysis, and EEG signal were submitted to a 0.1-30 Hz band-pass filter, with a sampling rate of 500 Hz. We then used independent component analysis to reduce ocular artifacts in EEGLAB v2023.1 ^50^. Following this, continuous recordings were epoched around feedback-locked from -200 to 1000 ms. Signals exceeding ±75μV in any given epoch were automatically excluded. Other artifacts were removed by visual inspection. For further statistical analysis, approximately 96.2% of the data were retained after excluding artifacts. Based on visual inspection of waveform plots and scalp topography distributions, and with reference to studies such as Jiang et al^51^ and Liu et al^24^, we selected a 220-320 ms time window with regions of interest (ROIs: F2, F1, Fz, FC2, FC1, FCz, C2, C1, Cz) for analyzing the feedback-related negativity (FRN) component, and a 300-450 ms time window with ROIs (FC1, FC2, FCz, C1, C2, Cz, CP1, CP2, CPz) for analyzing the P300 component.

### 4.7 Statistical analyses

#### 4.7.1 Behavioral data analysis

Firstly, generalized linear mixed models (GLMM) were conducted to examine the main effects and interactions of Context and Environmental volatility on behavioral metrics(R 4.0.1, lme4 package) ^52^. Two binomial GLMMs were constructed to focus on two behavioral metrics respectively: Accuracy and the probability of rule switching after receiving error feedback (i.e., *P*(RuleSwitch)). Here, accuracy was defined as the probability of selecting the optimal option in the current trial, i.e., the option corresponding to the high-probability association. For each model, Context and Environmental volatility were included as fixed effects, with reward conditions and stable conditions set as baselines, and participants were included as random effects in the model. Accuracy and *P*(RuleSwitch) were separately incorporated as dependent variables for model fitting.

Additionally, we conducted paired *t* tests to further analyze the differences in individual behavioral flexibility across the reward-punish conditions. We calculated the behavioral flexibility in Accuracy (ΔAccuracy, the difference between stable and volatile condition) and in the probability of switching rules after error feedback (Δ*P*(RuleSwitch)), the difference between stable and volatile condition) for each participant, separately for reward and punishment contexts. Paired *t* tests were then performed on these two difference variables to examine flexibility differences across contexts.

Finally, we tested whether the distinction in reward-punish learning also reflected in the model parameter. Paired *t* tests were used to test the asymmetry in reward-punish learning on learning parameters: learning rate, prediction error, and temperature parameter from decision model.

#### 4.7.2 EEG data analysis

To investigate the neural correlates of reward and punishment learning processes and their relationships with Hierarchical Gaussian Filter (HGF) model parameters, we employed linear mixed models (LMMs) to fit trial-by-trial EEG data with behavioral and computational predictors. Two separate LMMs were constructed to analyze the FRN and P300 components, respectively. The dependent variable in each model was the trial-by-trial EEG amplitude, averaged over specified time windows and regions of interest (ROIs). Context, Environmental volatility, and the absolute value after standardizing the precision weighted prediction error (|pwPE2|) were included as predictor variables, with participants modeled as random effects. The models were fitted using the restricted maximum likelihood estimation (REML) method to ensure parameter estimation stability. To evaluate the contribution of |pwPE2|, we compared two LMMs for each EEG component: a baseline model (including only Context, Environmental volatility, and their interaction) and a full model (additionally including |pwPE2| and its interactions). Model fit was assessed using a likelihood ratio test, where significant improvements indicated the importance of |pwPE2| in explaining EEG variance.

## 5 Limitation

The current study may have some limitations. First, although our sample size was sufficient to examine within-study effects, we may need to be more cautious in generalizing our results to larger populations. Therefore, subsequent studies can increase the sample size to obtain more stable effects and better generalize the results. In addition, although we used the same operation in the reward and punishment situations, these two situations may have different degrees of arousal to the subjects. Therefore, subsequent studies can increase the synchronous measurement of autonomic arousal, such as physiological indicators such as pupil diameter or heart rate.

## 6 Conclusion

This study indicates that FRN and P300 encode early significance detection and later value update, modulated by the precision-weighted prediction error (|pwPE2|) to reflect the processing of prediction errors at different hierarchical levels. We revealed differences in the two-stage processing across reward and punishment contexts: punishment contexts enhanced early salience detection (FRN), while late value updating (P300) was significant in both tasks but stronger in the punishment context. This disparity indicates that the motivational framework dynamically shapes the neural mechanisms of feedback processing by modulating processing priorities. The application of the HGF model reveals that |pwPE2| as the core driving signal, quantifying how the brain adjusts learning strategies based on environmental volatility and context. This study deepens our understanding of the neural mechanisms that underlie reward and punishment learning. Furthermore, it furnishes novel evidence that the two-stage processing theory applies in dynamic environments, where it is adaptively modulated by motivational contexts.

## Data availability

The data that support the findings of this study are available on request from the corresponding author.

## Code availability

The code of this study is available from the corresponding author upon request.

## Acknowledgements

We gratefully acknowledge the financial support from the National Natural Science Foundation of China (No. 32400872, 32400873). We also sincerely appreciate all the volunteers who participated in this study.

## Contributions

L.X. contributed to conceptualization, data curation, methodology, formal analysis, investigation, writing—original draft, and visualization. M.L. contributed to conceptualization, funding acquisition, writing—review and editing and supervision. W.L. contributed to funding acquisition, supervision, writing - review & editing, project administration.

## Competing interests

No potential conflict of interest was reported by the authors.

## Notes

### Competing Interest Statement

The authors have declared no competing interest.

## References

1. Combrisson, E. et al. Neural interactions in the human frontal cortex dissociate reward and punishment learning. eLife 12, RP92938 (2024).

2. Schultz, W., Dayan, P. & Montague, P. R. A neural substrate of prediction and reward. Science 275, 1593–9 (1997).

3. Schultz, W. Dopamine reward prediction-error signalling: a two-component response. Nat Rev Neurosci 17, 183–195 (2016).

4. Thorndike, L. & Bruce, D. Animal Intelligence: Experimental Studies. (Routledge, 2017).

5. Michely, J., Eldar, E., Erdman, A., Martin, I. M. & Dolan, R. J. Serotonin modulates asymmetric learning from reward and punishment in healthy human volunteers. Commun Biol 5, (2022).

6. Behrens, T. E. J., Woolrich, M. W., Walton, M. E. & Rushworth, M. F. S. Learning the value of information in an uncertain world. Nat. Neurosci. 10, 1214–1221 (2007).

7. Browning, M., Behrens, T. E., Jocham, G., O’Reilly, J. X. & Bishop, S. J. Anxious individuals have difficulty learning the causal statistics of aversive environments. Nat. Neurosci. 18, 590-+ (2015).

8. McCoy, B. & Lawson, R. P. Anxious individuals are more sensitive to changes in outcome variability and value differences in dynamic environments. Preprint at 10.1101/2024.08.25.609575 (2024).

9. Simoens, J., Verguts, T. & Braem, S. Learning environment-specific learning rates. PLoS Comput Biol 20, e1011978 (2024).

10. Bodi, N. et al. Reward-learning and the novelty-seeking personality: a between- and within-subjects study of the effects of dopamine agonists on young Parkinson’s patients. Brain 132, 2385–2395 (2009).

11. Elster, E. M. et al. Impaired Punishment Learning in Conduct Disorder. J. Am. Acad. Child Adolesc. Psychiatr. 63, 454–463 (2024).

12. Henco, L. et al. Aberrant computational mechanisms of social learning and decision-making in schizophrenia and borderline personality disorder. PLoS Comput Biol 16, e1008162 (2020).

13. Lawson, R. P., Mathys, C. & Rees, G. Adults with autism overestimate the volatility of the sensory environment. Nat Neurosci 20, 1293–1299 (2017).

14. Aylward, J. et al. Altered learning under uncertainty in unmedicated mood and anxiety disorders. Nat Hum Behav 3, 1116–1123 (2019).

15. Sharp, P. B., Russek, E. M., Huys, Q. J., Dolan, R. J. & Eldar, E. Humans perseverate on punishment avoidance goals in multigoal reinforcement learning. eLife 11, e74402 (2022).

16. Deng, G. et al. Dissociated modulations of intranasal vasopressin on prosocial learning between reward-seeking and punishment-avoidance. Psychol. Med. 53, 5415–5427 (2023).

17. Farashahi, S., Donahue, C. H., Hayden, B. Y., Lee, D. & Soltani, A. Flexible combination of reward information during choice under uncertainty. Nat Hum Behav 3, 1215–1224 (2019).

18. Mathys, C., Daunizeau, J., Friston, K. J. & Stephan, K. E. A Bayesian foundation for individual learning under uncertainty. Front. Hum. Neurosci. 5, 39 (2011).

19. Mathys, C. D. et al. Uncertainty in perception and the Hierarchical Gaussian Filter. Front. Hum. Neurosci. 8, (2014).

20. Fritsch, M., Weilnhammer, V., Thiele, P., Heinz, A. & Sterzer, P. Sensory and environmental uncertainty in perceptual decision-making. iScience 26, 106412 (2023).

21. Westwood, S. & Philiastides, M. G. Early Salience Signals Predict Interindividual Asymmetry in Decision Accuracy Across Rewarding and Punishing Contexts. Human Brain Mapping 45, e70072 (2024).

22. Hein, T. P., De Fockert, J. & Ruiz, M. H. State anxiety biases estimates of uncertainty and impairs reward learning in volatile environments. NeuroImage 224, 117424 (2021).

23. Iglesias, S. et al. Hierarchical prediction errors in midbrain and basal forebrain during sensory learning. Neuron 80, 519–530 (2013).

24. Liu, M., Dong, W., Qin, S., Verguts, T. & Chen, Q. Electrophysiological Signatures of Hierarchical Learning. Cerebral Cortex 32, 626–639 (2022).

25. Liu, M. et al. Modulating hierarchical learning by high-definition transcranial alternating current stimulation at theta frequency. Cerebral Cortex 33, 4421–4431 (2023).

26. Holroyd, C. B. & Coles, M. G. H. The neural basis of human error processing: Reinforcement learning, dopamine, and the error-related negativity. Psychological Review 109, 679–709 (2002).

27. Jollans, L. et al. Computational EEG modelling of decision making under ambiguity reveals spatio-temporal dynamics of outcome evaluation. Behavioural Brain Research 321, 28–35 (2017).

28. Xia, L., Xu, P., Yang, Z., Gu, R. & Zhang, D. Impaired probabilistic reversal learning in anxiety: Evidence from behavioral and ERP findings. NeuroImage-Clin. 31, 102751 (2021).

29. Fouragnan, E., Retzler, C., Mullinger, K. & Philiastides, M. G. Two spatiotemporally distinct value systems shape reward-based learning in the human brain. Nat Commun 6, 8107 (2015).

30. Walsh, M. M. & Anderson, J. R. Learning from experience: Event-related potential correlates of reward processing, neural adaptation, and behavioral choice. Neuroscience & Biobehavioral Reviews 36, 1870–1884 (2012).

31. Pfabigan, D. M., Alexopoulos, J., Bauer, H. & Sailer, U. Manipulation of feedback expectancy and valence induces negative and positive reward prediction error signals manifest in event-related brain potentials. Psychophysiology 48, 656–664 (2011).

32. Potts, G. F., Martin, L. E., Burton, P. & Montague, P. R. When Things Are Better or Worse than Expected: The Medial Frontal Cortex and the Allocation of Processing Resources. Journal of Cognitive Neuroscience 18, 1112–1119 (2006).

33. Polich, J. Updating p300: An integrative theory of P3a and P3b. Clin. Neurophysiol. 118, 2128–2148 (2007).

34. San Martín, R. Event-related potential studies of outcome processing and feedback-guided learning. Front. Hum. Neurosci. 6, (2012).

35. Nassar, M. R., Bruckner, R. & Frank, M. J. Statistical context dictates the relationship between feedback-related EEG signals and learning. eLife 8, e46975 (2019).

36. Yu, L. Q., Wilson, R. C. & Nassar, M. R. Adaptive learning is structure learning in time. Neurosci. Biobehav. Rev. 128, 270–281 (2021).

37. Cavanagh, J. F., Frank, M. J. & Allen, J. J. B. Social stress reactivity alters reward and punishment learning. Social Cognitive and Affective Neuroscience 6, 311–320 (2011).

38. Duek, O., Shahar, G., Osher, Y. & Kofman, O. Role of self-criticism in reward and punishment probabilistic learning. Personality and Individual Differences 134, 155–163 (2018).

39. Gueguen, M. C. M. et al. Anatomical dissociation of intracerebral signals for reward and punishment prediction errors in humans. Nat. Commun. 12, 3344 (2021).

40. Roth, A. M. et al. Punishment Leads to Greater Sensorimotor Learning But Less Movement Variability Compared to Reward. Neuroscience 540, 12–26 (2024).

41. Woody, E. Z. & Szechtman, H. Adaptation to potential threat: The evolution, neurobiology, and psychopathology of the security motivation system. Neuroscience & Biobehavioral Reviews 35, 1019–1033 (2011).

42. Galea, J. M., Mallia, E., Rothwell, J. & Diedrichsen, J. The dissociable effects of punishment and reward on motor learning. Nat Neurosci 18, 597–602 (2015).

43. McNaughton, N. & Corr, P. J. The neuropsychology of fear and anxiety: a foundation for Reinforcement Sensitivity Theory. in The Reinforcement Sensitivity Theory of Personality (ed. Corr, P. J.) 44–94 (Cambridge University Press, 2008).

44. Grubb, M. A. et al. The composition of the choice set modulates probability weighting in risky decisions. Cogn Affect Behav Neurosci 23, 666–677 (2023).

45. Kang, H. Sample size determination and power analysis using the G*Power software. J Educ Eval Health Prof 18, 17 (2021).

46. Faul, F., Erdfelder, E., Lang, A.-G. & Buchner, A. G*Power 3: A flexible statistical power analysis program for the social, behavioral, and biomedical sciences. Behav. Res. Methods 39, 175–191 (2007).

47. Kool, W., McGuire, J. T., Rosen, Z. B. & Botvinick, M. M. Decision Making and the Avoidance of Cognitive Demand. J. Exp. Psychol.-Gen. 139, 665–682 (2010).

48. Peirce, J. et al. PsychoPy2: Experiments in behavior made easy. Behav Res 51, 195–203 (2019).

49. Stephan, K. E., Penny, W. D., Daunizeau, J., Moran, R. J. & Friston, K. J. Bayesian model selection for group studies. NeuroImage 46, 1004–1017 (2009).

50. Delorme, A. & Makeig, S. EEGLAB: an open source toolbox for analysis of single-trial EEG dynamics including independent component analysis. Journal of Neuroscience Methods 134, 9–21 (2004).

51. Jiang, D. et al. Trait anxiety and probabilistic learning: Behavioral and electrophysiological findings. Biol. Psychol. 132, 17–26 (2018).

52. Bates, D., Mächler, M., Bolker, B. & Walker, S. Fitting Linear Mixed-Effects Models Using **lme4**. J. Stat. Soft. 67, (2015).

