## supplemental table for "Belief updating in uncertain environments are differentially sensitive to reward and punishment learning: Evidence from ERP"

**Supplementary Tables**

Table S1. Fixed Effects of the Accuracy Fitting Model

| | $\beta$(*S.E*) | *z* | *p* | 95%CI |
|---|---|---|---|---|
| (Intercept) | 1.72(0.07) | 24.31 | <0.001 | [1.59, 1.86] |
| Task type (punish) | -0.22(0.05) | -4.20 | <0.001 | [-0.32, -0.12] |
| Volatility (volatile) | -0.52(0.05) | -10.43 | <0.001 | [-0.62, -0.42] |
| Task type (punish): volatility (volatile) | 0.31(0.07) | 4.39 | <0.001 | [0.17, 0.44] |

Table S2. Fixed Effects of the P(RuleSwitch) Fitting Model

| | *β*(*S.E*) | *z* | *p* | 95%CI |
|---|---|---|---|---|
| (Intercept) | -2.09(0.08) | -27.08 | <0.001 | [-2.24, -1.94] |
| Task type (punish) | 0.3(0.06) | 5.11 | <0.001 | [0.18, 0.41] |
| Volatility (volatile) | 0.16(0.06) | 2.63 | 0.008 | [0.04, 0.27] |
| Task type (punish): volatility (volatile) | -0.23(0.08) | -2.82 | 0.005 | [-0.39, -0.07] |

Table S3. Comparison between models with and without parameters under FRN and P300 components.

| | | Fixed effect | Random effect | AIC | BIC |
|---|---|---|---|---|---|
| FRN | Model-full | z_\|pwPE2\| | Subject | 154518 | 154597 |
| | Model-reduced | | Subject | 154558 | 154606 |
| P300 | Model-full | z_\|pwPE2\| | Subject | 153044 | 153123 |
| | Model-reduced | | Subject | 153472 | 153520 |

Table S4. Fixed Effects of the FRN Model Without Parameter Modulation

| | $\beta(S.E)$ | $t$ | $p$ | 95%CI |
|---|---|---|---|---|
| (Intercept) | 5.57(1.13) | 4.94 | <0.001 | [3.36, 7.78] |
| Task type (punish) | -0.00(0.21) | -0.02 | 0.98 | [-0.41, 0.40] |
| Volatility (volatile) | -0.75(0.21) | -3.64 | <0.001 | [-1.15, -0.35] |
| Task type (punish): volatility (volatile) | -0.37(0.29) | -1.27 | 0.21 | [-0.94, 0.20] |

Table S5. Fixed Effects of the P300 Model Without Parameter Modulation

| | $\beta$(*S.E*) | *t* | *p* | 95%CI |
|---|---|---|---|---|
| (Intercept) | 7.02(0.91) | 7.73 | <0.001 | [5.24, 8.80] |
| Task type (punish) | -0.16(0.20) | -0.80 | 0.42 | [-0.55, 0.23] |
| Volatility (volatile) | -1.12(0.20) | -5.57 | <0.001 | [-1.51, -0.72] |
| Task type (punish): volatility (volatile) | -0.31(0.28) | -1.10 | 0.27 | [-0.87, 0.25] |

Table S6. Fixed Effects of the FRN Model with Parameter Modulation

| | $\beta$(*S.E*) | *t* | *p* | 95%CI |
|---|---|---|---|---|
| (Intercept) | 5.57(1.12) | 4.95 | <0.001 | [3.36, 7.77] |
| Task type (punish) | 0.01(0.21) | 0.06 | 0.95 | [-0.39, 0.42] |
| Volatility (volatile) | -0.74(0.21) | -3.57 | <0.001 | [-1.14, -0.33] |
| $\lvert pwPE_2 \rvert$ | -0.06(0.16) | -0.35 | 0.73 | [-0.37, 0.26] |
| Task type (punish): volatility (volatile) | -0.49(0.30) | -1.67 | 0.09 | [-1.06, 0.08] |
| Task type (punish): $\lvert pwPE_2 \rvert$ | 0.90(0.21) | 4.38 | <0.001 | [0.50, 1.30] |
| Volatility (volatile): $\lvert pwPE_2 \rvert$ | 0.15(0.22) | 0.69 | 0.49 | [-0.28, 0.59] |
| Task type (punish): volatility (volatile): $\lvert pwPE_2 \rvert$ | -0.40(0.30) | -1.34 | 0.18 | [-0.97, 0.18] |

Table S7. Fixed Effects of the P300 Model with Parameter Modulation

| | *β*(*S.E*) | *t* | *p* | 95%CI |
|---|---|---|---|---|
| (Intercept) | 7.18(0.91) | 7.93 | <0.001 | [5.40, |
| Task type (punish) | -0.29(0.20) | -1.48 | 0.138 | [-0.68, 0.10] |
| Volatility (volatile) | -1.21(0.20) | -6.09 | <0.001 | [-1.60, -0.82] |
| $\|pwPE_2\|$ | 1.50(0.15) | 9.81 | <0.001 | [1.20, 1.80] |
| Task type (punish): volatility (volatile) | -0.51(0.28) | -1.82 | 0.069 | [-1.07, 0.04] |
| Task type (punish): $\|pwPE_2\|$ | 0.54(0.20) | 2.73 | 0.006 | [0.15, 0.93] |
| Volatility (volatile): $\|pwPE_2\|$ | -0.02(0.21) | -0.08 | 0.935 | [-0.44, 0.40] |
| Task type (punish): volatility (volatile): $\|pwPE_2\|$ | -0.35(0.29) | -1.23 | 0.218 | [-0.91, 0.21] |
